# miR-551a and miR-551b-3p target GLIPR2 and promote tumor growth in High-Risk Head and Neck Cancer by modulating autophagy

**DOI:** 10.1101/019406

**Authors:** Narasimha Kumar Karanam, Liang-Hao Ding, Tae Hyun Hwang, Hao Tang, Michael D. Story

## Abstract

Distant metastasis (DM) and local-regional recurrence (LR) after radiation and chemo therapy are major cause of treatment failure for patients with head and neck squamous cell carcinoma. However, detailed underlying mechanisms leading to DM and LR in patients are not fully understood yet. MiRNA have been proposed as biomarkers in a variety of biological and medical conditions such as cancer and stress response. The advantages of miRNA as a biomarker lies in its stability in tissues as well as body fluids, hence the potential for non-invasive diagnosis and prognosis. In this study, towards understanding the molecular mechanism causing DM and LR in HN cancer patients we performed miRNA expression profiling using tumor samples from 118 head and neck cancer patients treated by post-operative radiotherapy (PORT) at M.D. Anderson Cancer Center from 1992 to 1999. All patients were considered to be at high-risk for recurrence having histologically proven advanced squamous cell carcinoma. Amongst these samples, 41 found to have distant metastasis (DM), 53 responded without relapse (no evidence of disease (NED)) to PORT. Comparison of miRNA expression between DM and NED specimens using two-way ANOVA identified 28 miRNAs that were differentially expressed with statistical significance (FDR < 0.2 and fold change > 1.5). Amongst these 28 miRNAs seen in the DM and NED outcome groups, miRNAs 551a and 551b are significantly associated with the DM group. Interestingly these two miRNAs share same seed sequence. Moreover Kaplan-Meir survival analysis in our data set and two other data sets suggested that miR-551a and miR-551b expressions are associated with poor survival in patients. We further performed cell proliferation, migration and invasion assays using the HN5 and UMSCC-17B head and neck cancer cell lines by transfection of either mimic or an inhibitor of miR-551a and miR-551b. The results suggested that miR-551a and miR-551b mimics promote proliferation, migration and invasion whereas the inhibitor decreased. Further studies indicated that these miRNAs target GLIPR2 expression and miR-551a, miR-551b and GLIPR2 axis at least in part plays an important role in tumor progression. Hence we need to further explore miR-551a and miR-551b-3p role in HN cancer progression in detail in in-vivo models to use them as therapeutic targets in future.

## Introduction

Head and neck squamous cell carcinoma (HNSCC) is the sixth leading malignancy diagnosed and the eighth most common cause of cancer death worldwide (Parfenov et al., 2014). Head and neck cancer consists of complex anatomical diverse groups of cancers and each of them demands different multi-modality management. Due to its complicated patterns of tumor invasion and occurrence in most sensitive parts of body which can impact vital functions in human body HNSCC are associated with high incidence of distant metastasis and local relapse resulting in high mortality rate. Latest advances in surgical, medical therapies including chemo and radio therapies for HNSCC have resulted in only 50% improvement in terms of mortality rate in the last three decades (Leon et al., 2001; Posner et al., 2007; Vermorken et al., 2007). This study includes 118 number of HNSCC patients who were treated by post-operative radiotherapy (PORT) at MD Anderson Cancer Center from 1992 to 1999. Moreover all these patients were considered to be at high-risk for recurrence/ metastasis having histologically proven advanced squamous cell carcinoma. Hence determining the necessity of systemic therapy and subjecting patients to the debilitating effects associated with such regimes has led to the pursuit of molecular-based markers for radio-chemo-sensitivity and the prediction of therapeutic outcomes.

After completion of human genome sequencing it was rather surprising to learn that only less than 2% of the human genome is protein coding (Lander et al., 2001; Venter et al., 2001). However later it was revealed that 65% of the genome is actually transcribed but not all translated into proteins which suggest that there is substantial proportion of non-coding transcripts/RNAs which are thought to be playing important role in gene expression regulation typically via complementarity to the 3’ UTR of target mRNA (Carninci et al., 2005). Micro RNAs belongs to the non-coding RNA group which are short stretch of (19 to 25 nucleotides) RNA molecules that are capable of inducing post-transcriptional silencing of its target mRNAs (Morris and Mattick, 2014). However some studies demonstrate that miRNA can also target the 5’ UTR of a target mRNA both open reading frames (ORF) and promoter regions (Lee et al., 2003; Place et al., 2008). First evidence of miRNAs existence was provided by Ambros and colleagues in 1993 in C. elgans (Lee et al., 1993). Later it was found that these miRNAs are evolutionarily conserved and many are tissue and lineage specific (Berezikov et al., 2006; Heimberg et al., 2010). Understanding miRNA regulatory networks is quite complex as in some instances single miRNA can regulate more number of target mRNAs (Fabian et al., 2010) and, reciprocally single mRNA could be targeted by a large number of miRNAs (Schnall-Levin et al., 2011), although the implied regulatory logic of this complex multiplex arrangement has not been explained. mRNAs are general targets of miRNAs but may also include other types of RNAs as well (Hansen et al., 2011). Functionally miRNAs have been shown to have regulatory role in many physiological, developmental and disease processes, including pluripotency (Leonardo et al., 2012), epithelial–mesenchymal transition and metastasis (Bracken et al., 2009), testis differentiation (Rakoczy et al., 2013), diabetes (Fernandez-Valverde et al., 2011), and neural plasticity and memory (Bredy et al., 2011), among others (Park et al., 2010). Interestingly it was found that human miRNA genes are frequently located at fragile sites and genomic regions involved in cancers (Calin et al., 2004). Alterations in miRNA expression have been observed in various solid tumors, including hepatocellular carcinoma (Gramantieri et al., 2008; Negrini et al., 2011) and lung (Lin et al., 2010), breast (O'day and Lal, 2010), pancreatic (Roldo et al., 2006), gastric (Chen et al., 2012) and prostate cancer (Porkka et al., 2007), and could potentially be used for cancer diagnosis, prognosis and response to treatment (Madhavan et al., 2013; Wang et al., 2012). Moreover miRNAs are highly stable due to their encapsulation in exosomes and association with protein complexes in bio-fluids such as serum or plasma, urine and saliva (Chen et al., 2008; Mitchell et al., 2008; Park et al., 2009), thus miRNAs can be used as non-invasive diagnostic and prognostic markers. With these advantages of miRNA research we have done extensive miRNA array profiling of HNSCC tumor samples.

Earlier there were some studies about miRNA profiling of HN tumor samples which include mostly primary tumors. But in this study we have done comprehensive miRNA and gene expression profiling of ∼100 HN tumor samples which are mostly secondary tumors. Moreover we further explored the important and novel role of miR-551a and miR-551b-3p in HN tumor progression and aggressiveness in-vitro which is in accordance with survival data of patients’ in-vivo.

## Materials and methods

**Tumor samples:** This study includes 118 number of HNSCC patients who were treated by post-operative radiotherapy (PORT) at MD Anderson Cancer Center from 1992 to 1999.

**Cell culture:** HN5 and UMSCC17B cell lines were established from Head and neck cancer patients are procured from MD Anderson cancer center and University of Michigan Medical School respectively. HN cancer cell lines were grown in Dulbecco’s Modified Eagle’s Medium (DMEM) (Sigma-Aldrich Co, St Louis, MO) containing high glucose (4500mg/L), 2mM L-glutamine, 1% nonessential amino acids, 1% Penicillin-Streptomycin (Invitrogen, Carlsbad, CA) and 10% fetal bovine serum in a humidified atmosphere of 5% CO_2_ at 37°C.

**RNA extraction:** Total RNA was extracted according to manufacturer’s protocol using Qiagen RNeasy Plus kit (Qiagen GMBH, Hilden, Germany) which retains small RNA fractions.

**miRNA array:** miRNA array profiling was done using Exiqon platform using modified manufacturer’s protocol. Briefly, 750 ng of RNA was labeled using Exiqon miRCURY LNA microRNA Hi-Power labeling kit labeling kit-Hy3 (Exiqon Inc, MA) labeling with miRCURY LNA array spike-in microRNA kit V2 (Exiqon Inc, MA). Then labeled cRNA was hybridized on miRCURY LNA microRNA array 7^th^ generation (Exiqon Inc, MA) slide using Agilent chambers (Agilent Technologies) at 56°C overnight. Next day morning these hybridzed slides were washed using exiqon buffers and finally scanned using Tecan’s Power Scanner (Tecan US Inc, Morrisville, NC) and features were extracted using Arrayit ImaGene 9 (Arrayit, CA) software for further analysis.

**Microarray:** Microarray profiling was done using Illumina HT12 V4 Beadchip platform using modified manufacturer’s protocol. 500ng of RNA was labeled using Illumina Total Prep kit (Invitrogen, TX). 750ng of this labeled cRNA was hybridized on Illumina HT12 BeadChip V-4 (Illumina, CA) using standard Illumina hybridization protocol and scanned using HiScan array scanner (Illumina, CA). Microarray data were background-subtracted and normalized using the MBCB algorithm.

**Quantitative RT-PCR:** 20 ng of RNA extracted from tissues was used as qRT-PCR reactions starting material. RNA was reverse transcribed using the miRCURY Locked Nucleic Acid (LNA^TM^)Universal Reverse Transcription (RT) microRNA PCR, Polyadenylation and cDNA synthesis kit (Exiqon Inc, MA, USA). Then, cDNA was diluted 50x and assayed in duplicates of 10 μl PCR reactions according to the manufacturers protocol using miR-551a and miR-551b-3p specific miRCURY LNA primers (Exiqon Inc, MA, USA) and U6 miRCURY LNA primer was used as loading control. qRT-PCR reaction was run on Peltier Thermal Cycler machine and ΔCT values were calculated after normalization using U6 expression values in each sample and relative quantification values were plotted using GraphPad Prism software (GraphPad Prism software Inc, CA, USA).

**Transfections:** We have used primarily two transfection reagents in this study i.e. 1) INTERFERin (Polyplus, Illkirch, France) and 2) Lipofectamine 2000 (Life Technologies, CA, USA). For miRNA mimics and inhibitors transfections we used IINTERFERin reagent and transfections were carried out according to the manufacturer’s instructions and taken care that cell density is about 40 to 50% at the time of transfection. Generally we have used 40 nM final concentrations of miRNA mimics and inhibitors for transfection unless otherwise specifically mentioned. For cDNA transfections we used Lipofectamine 2000 reagent and transfections were carried out according to manufactures’ instructions and taken care that cell density is about 80 to 90% at the time of transfection. Approximately 1μg cDNA per 1 million cells was used for transfection.

**CellTiter Glo cell viability assays:** HN5 and UMSCC-17B cells were grown to 40 to 50% confluence on the day of transfection in 96 well plates. These two cell lines were transfected with miR-551a and miR-551b-3p mimics (Life Technologies)and inhibitors and GLIPR2 siRNA (Thermo Scientific) at 40 nM final concentration using Interferin transfection reagent (Polyplus, Illkirch, France) according to manufacturer’s protocol. After transfection with indicated materials cells were grown for indicated time points as shown in figures. On the day of doing proliferation assay 25 ul of CTG assay buffer (Promega) was added to each well which quantitates ATP production produced by metabolically active cells and this luminescence signal intensities were read by 96 well plate reader (PolarStar Optima, BMG LabTech, NC, USA). For quantification purpose relative luminescence unit values (RLU) were plotted for each cell line and time point.

**Migrations assays:** Cell migration assay was carried out according to modified protocol from Wiggins HL et al. In brief-∼1% low melting agarose was prepared in PBS and heated in 100°C water bath for 15 min and 6 micro liters of 3 agarose spots were spotted in 12 well plate and allowed them to solidify for 15 min under UV light in hood. Meanwhile cells were harvested and counted. About 200 thousand cells were seeded gently in wells containing agarose spots. Next day morning cell culture media was changed and transfections with indicated reagents and materials were carried out. After 48 hrs agarose spots were removed and fresh media with various drugs at given concentration was added and let the cells to migrate for 72 hrs. at the end of 72 hrs media was removed and cells were stained with crystal violet solution and images were taken using Evos XL microscope (Electron Microscopy Sciences, PA) at 4X magnification. Here in the below picture agarose spot at day1 was shown and all other spots were stained at day 5 from seeding. Statistical significance was calculated by using Students’ *t* test.

**Invasion assays:** HN5 and UMSCC-17B cells were grown in 6 well plate up to 40 to 50% conluency and then transfected with materials and reagents indicated in figures at desired concentration and grown for 2 days. Cells were serum starved overnight after 36 hours of transfection using serum free DMEM media containing all other supplements. Next day morning BD matrigel invasion chambers were rehydrated using serum free DMEM medium in normal cell culture incubators for 2 hrs. After rehydration of matrigel wells transfected cells which are serum starved for overnight were harvested and counted. Approximately 100 K cells in serum free DMEM medium were loaded in upper chamber of well and 5% FBS containing DMEM medium was loaded in lower chamber of well which serves as chemoattractant. Cells were allowed to invade through matrigel for 24 hrs. Next day the inner sides of wells containing matrigel together with non-invaded cells were cleaned using cotton swab. The lower side of well containing invaded cells was fixed using 70% ethanol and stained using crystal violet solution (recipe). After staining images of cells were taken using Evos XL microscope (Electron Microscopy Sciences, PA) at 10X magnification at 4 different fields and number of cells per field were counted and graphed. Statistical significance was calculated by using Students’ *t* test.

**Luciferase reporter assay:** The 3’ UTR sequence of GLIPR2 which was predicted to have complementarities with miRNAs was cloned into pmiRGLO vector (Promega) using Sac I and Xho I restriction enzymes. Later on cloning was validated by sequencing the product. Cloned 3’UTR GLIPR2 plasmid was used for Luciferase reporter assay.

**Cell lysate preparation and Westernblots:** Western blot analysis was conducted using whole-cell lysates. After 3 days of transfection, the transfected cells were harvested for protein extraction. Total cell lysates from different experiments were obtained using CellLytic M buffer (Sigma Aldrich Co, Mo, USA) by lysing the cells in the protein lysis buffer from (Thermo Scientific) containing protease inhibitor cocktail and phosphatase inhibitor. Protein estimation of these cell lysates was done using Bradford’s assay as per manufacturer’s instructions and 25 μg of protein was loaded on 12% SDS-PAGE gel and western blot analysis was conducted as previously described elsewhere (Karanam et al., 2010). We used LC3, p62, AKT,phosphor AKT (S-473), phosphor p38 MAPK, p27 Kip1 (Cell Signaling, MA, USA) and Actin (Sigma Aldrich Co, MO, USA). GLIPR2 antibody was a kind gift from Dr. Beth Levine laboratory.

### Statistics and KM curves analysis

Differences in miRNA expression among patients or sample groups were analysed using Anova statistical analysis. Bioinformatics analysis of predicted pathways influenced by the panel of selected miRNAs was investigated using the Diana miRPath online tool (http://diana.cslab.ece.ntua.gr/pathways/index_multiple.php) using three different algorithms (DIANA-microT v4.0, TargetScan 5 and PicTar 4-way). Kaplan-Meier survival curve was generated using R package “survival”.

## Results

### Head and neck tumor samples miRNA profiling revealed higher expression of miR-551b-3p in DM patients compared to NED patients

miRNA profiling of HN tumor samples was carried out on Illumina platform as described in materials and methods section. miRNA array analysis revealed that xx number of miRNAs were upregulated and xx number of miRNAs were downregulated. Among these differentially expressed miRNAs we miR-551b-3p was upregulated ∼1.5 foled in DM patients compared to NED patients with p-value of 0.04 (Fig1.a). We studied about miR-551b-3p function further as there were not many reports so far about miR-551b-3p role in cancer and non in HN cancer especially. Besides miR-551b-3p we looked for miR-551a expression as well in these samples as it contains same seed sequence as of miR-551b-3p. Interestingly miR-551a was also upregulated ∼1.2 fold with p value 0.04 (Fig1.a). We hypothesized that these two miRNA potentially target same target as these two miRNAs share same seed sequence.

**Fig.1:**
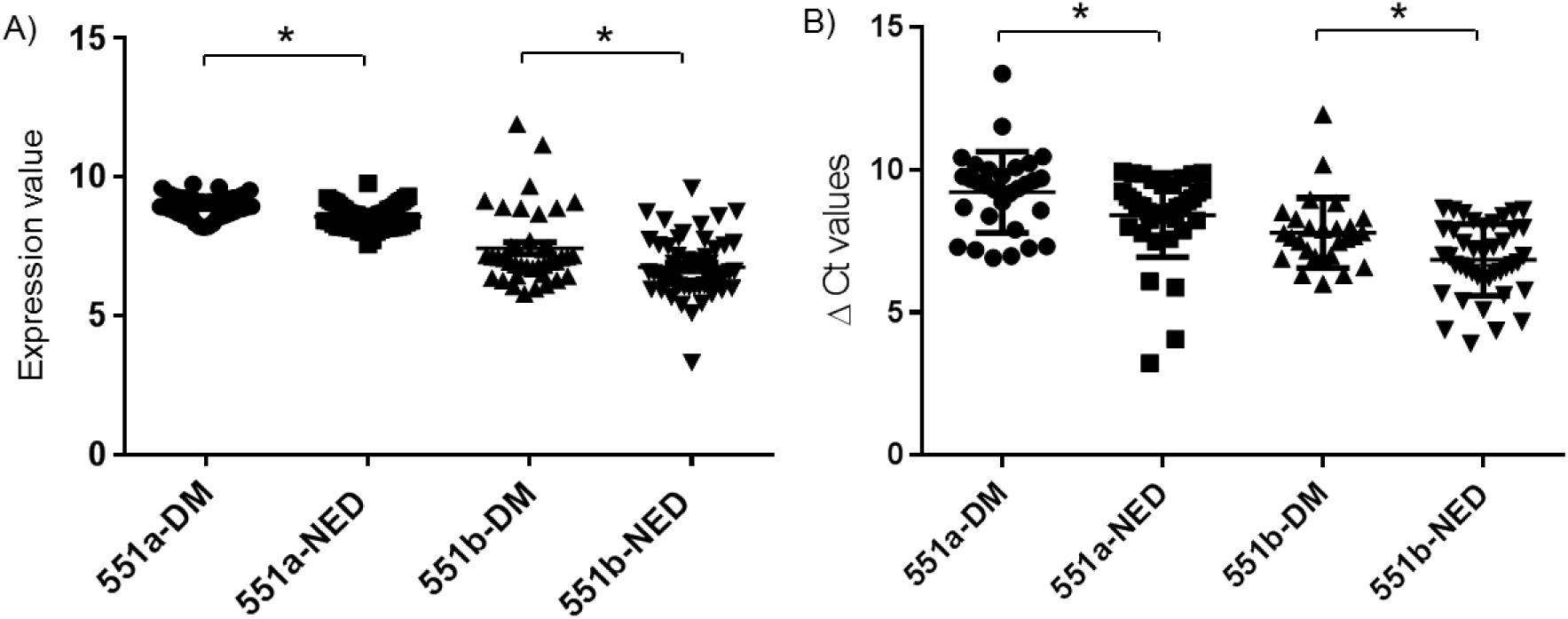
miR-551a and miR-51b-3p gene expression levels in tissue samples and validation of their over expression by qRT-PCR. A) Normalized expression values of miR-551a and miR-551b-3p from Exiqon miRCURY LNA microRNA array in DM and NED groups of HN cancer patients. B) qRT-PCR validation of over expression of miR-551a and miR-551b-3p in DM HN patients’ samples compared to NED HN patients’ samples using respective specific Exiqon pimers normalized using U6 miRNA expression values. Statistically significant (p-value > 0.05) differences were denoted by asterisks on top of the groups compared.

Later on we validated our miRNA array observation of upregulation of miR-551a and miR-551b-3p using specific Exiqon LNA primers and we used U6 as internal control for standardization purpose. We found that miR-551a and miR-551b-3p are indeed upregulated in DM patients compared to NEDs by qRT-PCR (Fig1.b).

### miR-551a and miR-551b-3p expressions correlate with HN patients survival

After having confirmation of miR-551a and mmiR-551b-3p higher level of expression in DM patients we were curious to know about its correlation with survival of patients which is very important clinical feature to evaluate. Interestingly we observed that in our data set both miR-551a and miR-551b-3p expression levels correlated negatively with patients’ survival i.e. patients with high level of miR-551a and miR-551b-3p expression have lower survival rate and vice versa (Fig2.a & b). In addition to our data set (GSE36682) we also found the same negative correlation in other HN cancer (Nasopharyngeal carcinoma) data set (Fig2.c & d). This critical observation led us to study in depth about miR-551a and miR-551b-3p molecular function in HN cancer cells.

**Fig.2:**
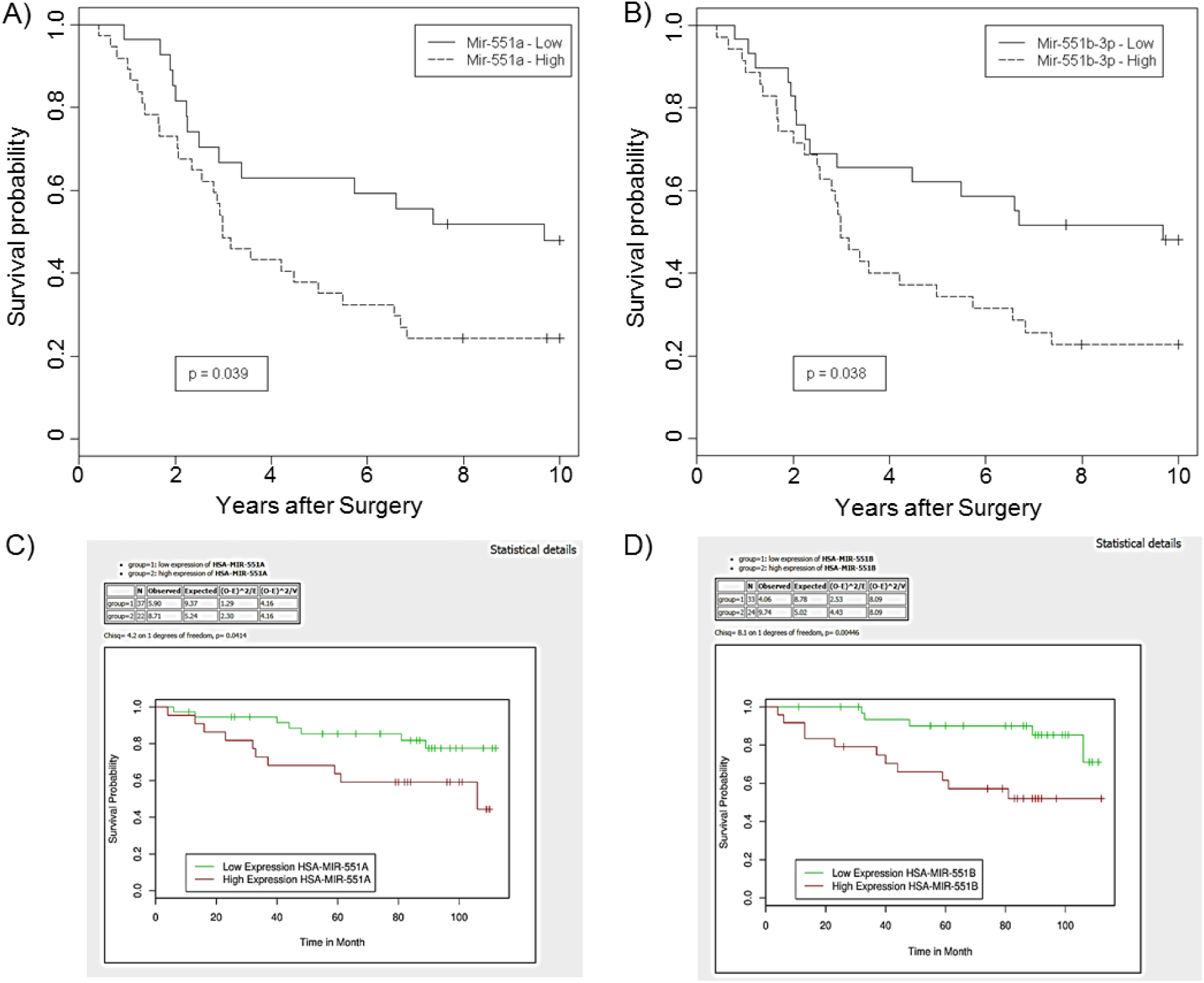
miR-551a and miR-551b-3p expression correlates with HN cancer patients survival data. KM curves were drawn using 10-year survival data available for HN cancer patients with A) miR-551a and B) miR-551b-3p expression data and it shows that the patients with higher miR-551a and miR-551b-3p expression values have lower survival and vice-versa. KM survival curves from GSE36682 nasopharyngeal carcinoma dataset for C) miR-551a and D) miR-551b-3p. p-values are mentioned in the graph.

### Ectopic expression of miR-551a and miR-551b-3p increases HN cancer cells proliferation, migration and invasion

Proliferation, migration and invasion are major contributors of tumor progression and metastatic behavior. We studied the functional role of miR-551a and miR-551b-3p in HN cancer cell lines-HN5 and UMSCC-17B which are established from patients by modulating miR-551a and miR-551b-3p expressions using synthetic mimics and inhibitors. When we transfected the HN cancer cell lines with miR-551a and miR-551b-3p mimics we observed increase in cell proliferation where as their inhibitors decreased the cell proliferation indicating miR-551a and miR-551b-3p role in cell proliferation (Fig3).

**Fig.3:**
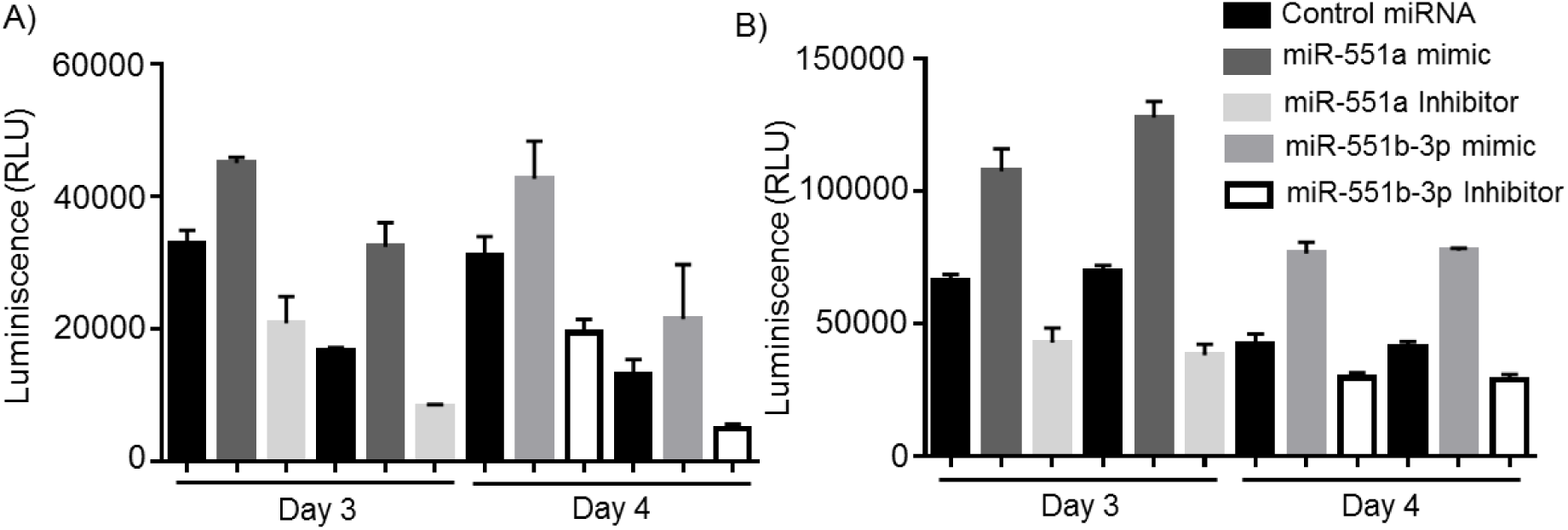
Modulation of miR-551a and miR-51b-3p levels using their mimics and inhibitors affect cell proliferation in HN5 cells. Effect of A) miR-551a and B) miR-551b-3p expression on cell proliferation was temporally monitored after transfection of their synthetic mimics and inhibitors in HN5 cells using CTG assay as described in methods. Ectopic expression of A) miR-551a and B) miR-551b-3p using their mimics increased the proliferation whereas decreased proliferation was observed after transfection of A) miR-551a and B) miR-551b-3p inhibitors in HN5 cells and the magnitude of change in proliferation was high on day 4 after transfection.

Wound healing assay using miR-551a and miR-551b-3p mimics increased the migration of HN5 and UMSCC-17B cell lines whereas miR-551a and miR-551b-3p inhibitors decreased (Fig4). We quantified the unclosed area from 2 independent experiments containing 4 circles in each experiment by measuring the diameter and graphed it as shown in figure 4.

**Fig.4:**
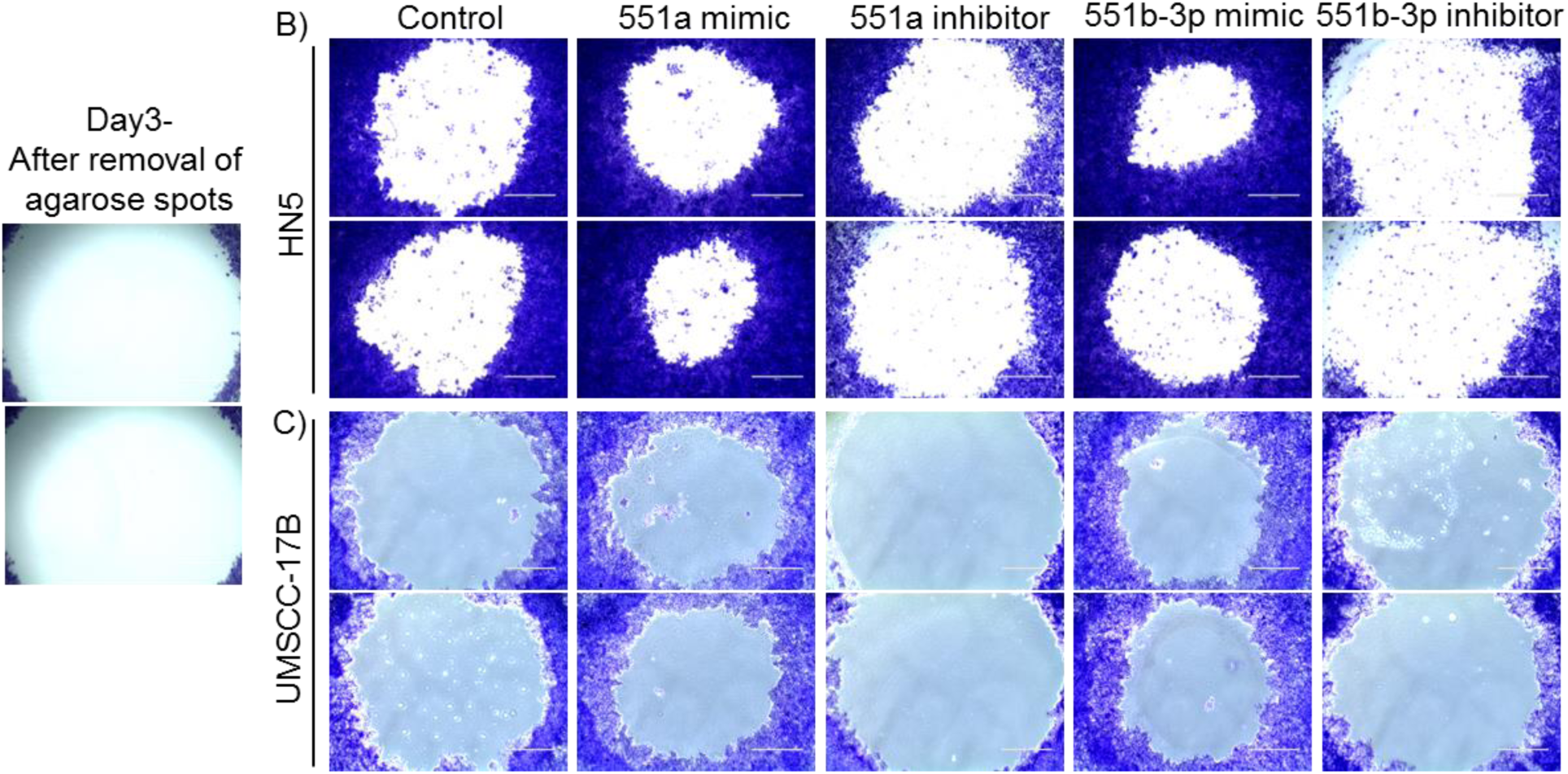
miR-551a and miR-551b-3p increase cell migration in HN5 cells. A) Ectopic expression of miR-551a and miR-551b-3p using their synthetic mimics increase cell migration where as their antagonistic inhibitors decrease cell migration in HN5 cells. After transfection of miR-551a and miR-551b-3p mimics and inhibitors equal area of agarose spots were removed and the area of open space left was calculated. Representative pictures of agarose spots at different stages and conditions were shown here. B) Area of open space left was calculated and represented by graph. Statistically significant values are denoted by asterisks which are less than p-value of 0.05.

Invasion assay was carried out using BD matrigel chambers and we used HT-1080 highly metastatic fibro sarcoma cells as positive control. MiR-551a and miR-551b-3p mimics increased the invasion capability of HN5 and UMSCC-17B cells whereas miR-551a and miR-551b-3p inhibitors decreased (Fig5). Number of invaded cells was counted from at least 4 sections of a chamber from experiment and average number of invaded cell per field was calculated and kept in graph as shown in Figure 5.

**Fig.5:**
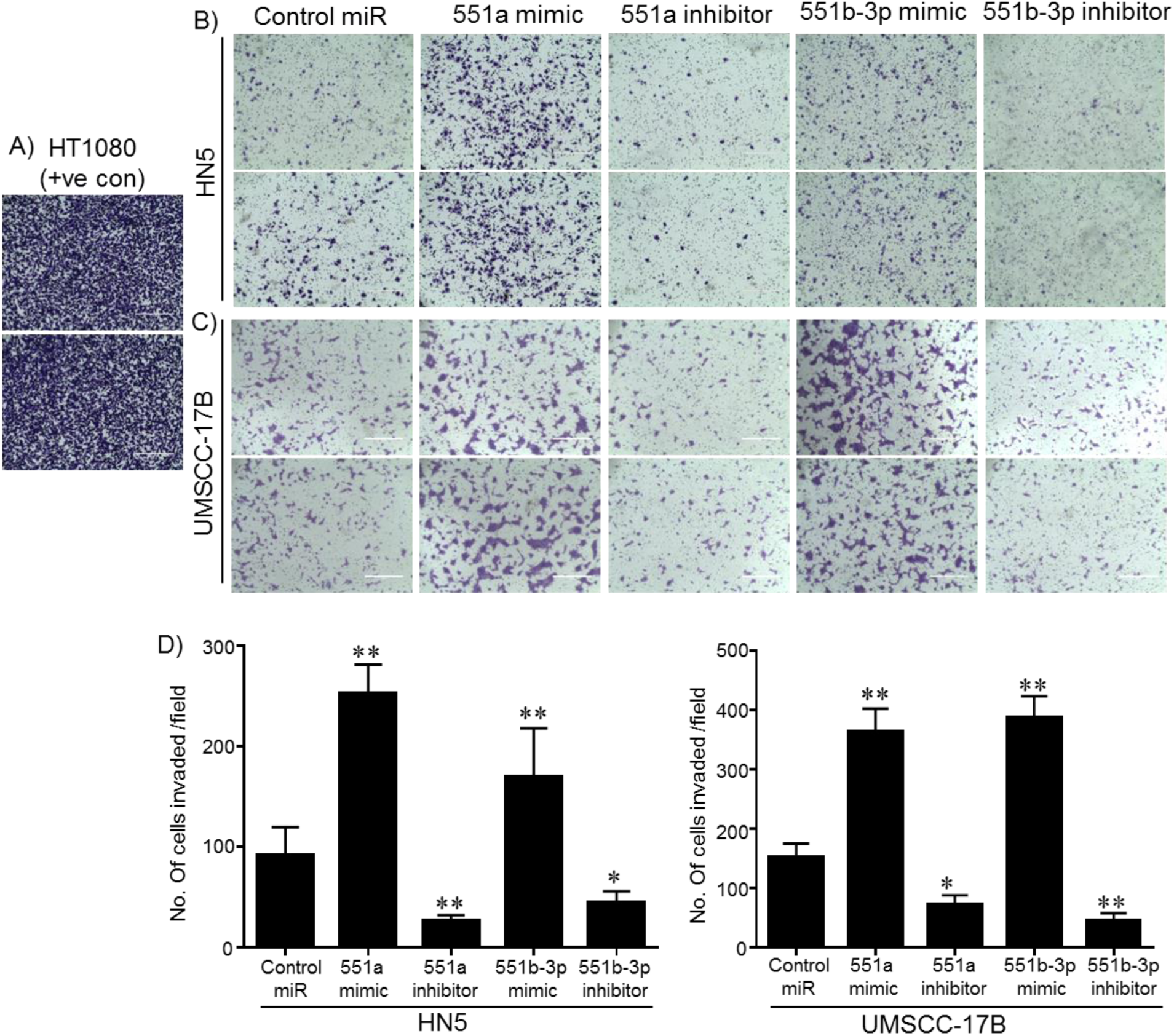
miR-551a and miR-551b-3p increases cell invasion ability in HN5 cells. A) HT-1080 fibrosarcoma cells, which are highly invasive in nature were used as positive control for invasion experiments. Ectopic expression of B) miR-551a C) miR-551b-3p using their synthetic mimics increase cell invasion ability where as their antagonistic inhibitors of B) miR-551a C) miR-551b-3p decrease cell invasion in HN5 cells. Two days after transfection of miR-551a and miR-551b-3p mimics and inhibitors equal number of transfected cells were transferred into invasion chambers where upper chamber contains no serum and lower camber contains 5% serum to form gradient to facilitate cell invasion. D) number of of invaded cells were calculated and represented by graph. Statistically significant values are denoted by asterisks which are less than p-value of 0.05.

### Micro array and bioinformatics analysis suggest GLIPR2 as potential target of miR-551a and miR-551b-3p

We have carried out micro array analysis to evaluate the gene expression changes upon transfection of miR-551a and miR-551b-3p mimics and inhibitors together with controls into HN5 cells to know potential targets of miR-551a and miR-551b-3p. We found out that GLIPR2 gene expression was decreased upon miR-551a and miR-551b-3p mimics transfection whereas miR-551a and miR-551b-3p inhibitors increased GLIPR2 gene expression compared to their respective controls (Fig 6. A). This data gave us a hint that GLIPR2 could be a potential target of miR-551a and miR-551b-3p. Later on we also found that miR-551a and miR-551b-3p contains complimentary sequence to 3’ UTR region of GLIPR2 (Fig 6.B). Together with these findings we also have data in HN tumors gene expression micro array analysis that GLIPR2 gene expression was low in DM patients compared to NEDs (Fig 6.C) which cumulatively suggest that GLIPR2 is a potential direct target of miR-551a and miR-551b-3p.

**Fig.6:**
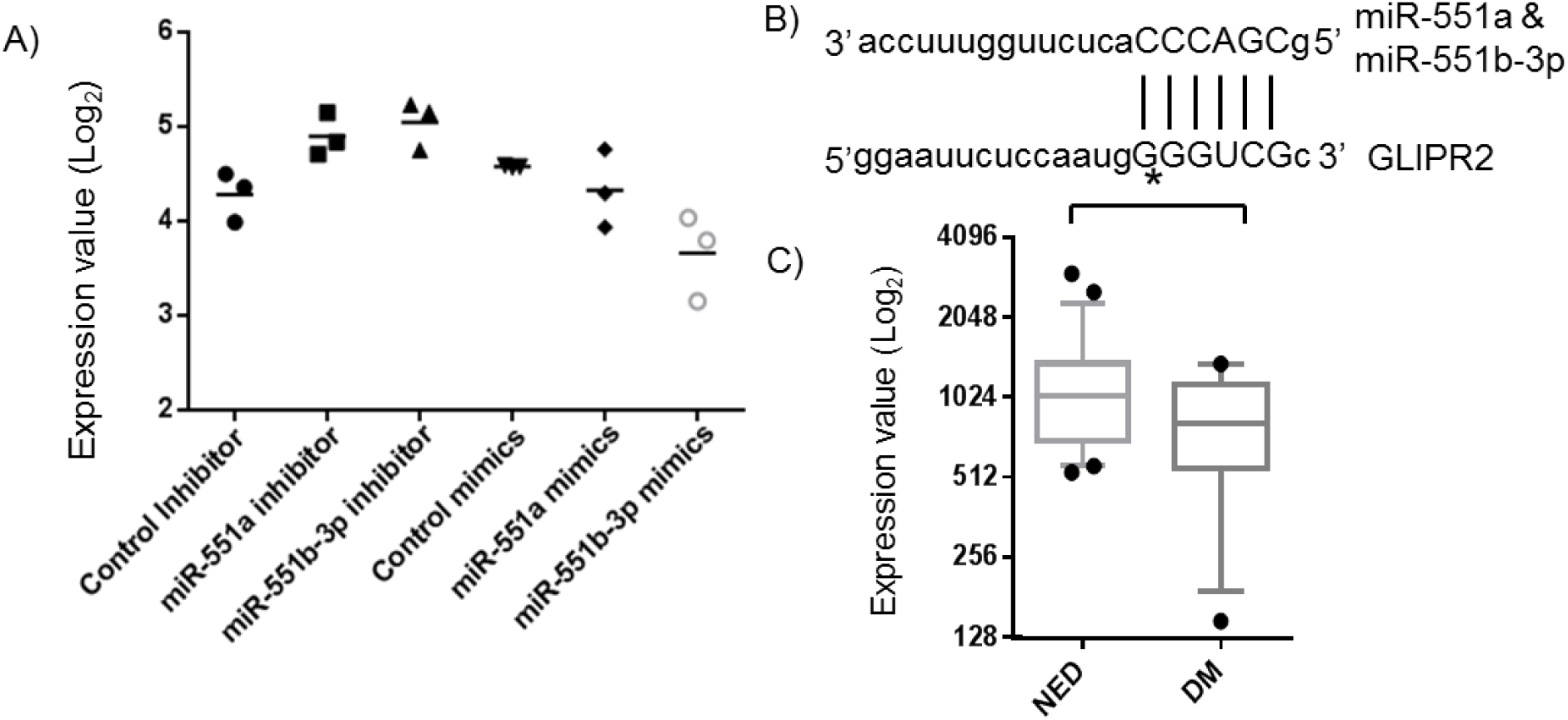
miR-551a and miR-551b-3p contains complimentary sequence in 3’ UTR region of GLIPR2. A) Alignment of miR-551a, miR-551b-3p seed sequence with 3’ UTR region of GLIPR2. B) Gene expression values (log2) of GLIPR2 in NED and DM groups of HN patients, which are inversely correlated with miR-551a, miR-551b-3p expression. C) Normalized microarray expression values in miR-551a and miR-551b-3p mimics and inhibitors treated HN5 cells compared to controls showing lower expression in mimic treated condition and higher expression in inhibitor treated condition compared to their respective controls. Statistically significant values are denoted by asterisks which are less than p-value of 0.05.

### Luciferase construct of GLIPR2 3’ UTR and western blot experiment confirms GLIPR2 is direct target of miR-551a and miR-551b-3p

Luciferase construct of GLIPR2 3’UTR (xx sequence region) was used to see direct physical interaction between GLIPR2 3’ UTR and miR-551a and miR-551b-3p. We observed clear repression in luciferase signal when we combined pmirGLO-GLIPR2 3’ UTR plasmid and miR-551a and miR-551b-3p mimics whereas control pmirGLO vector alone didn’t have such effect suggesting a physical interaction between miR-551a and miR-551b-3p and GLIPR2 3’ UTR (Fig 7.A). After confirmation using luciferase construct of GLIPR2, we used GLIPR2 specific antibody to see whether there is any GLIPR2 protein level change upon transfection of miR-551a and miR-551b-3p mimics and inhibitors. Using GLIPR2 antibody we found that there is clear reduction in protein levels of GLIPR2 upon miR-551a and miR-551b-3p mimics transfection but not much difference observed upon miR-551a and miR-551b-3p inhibitors transfection.

**Fig.7:**
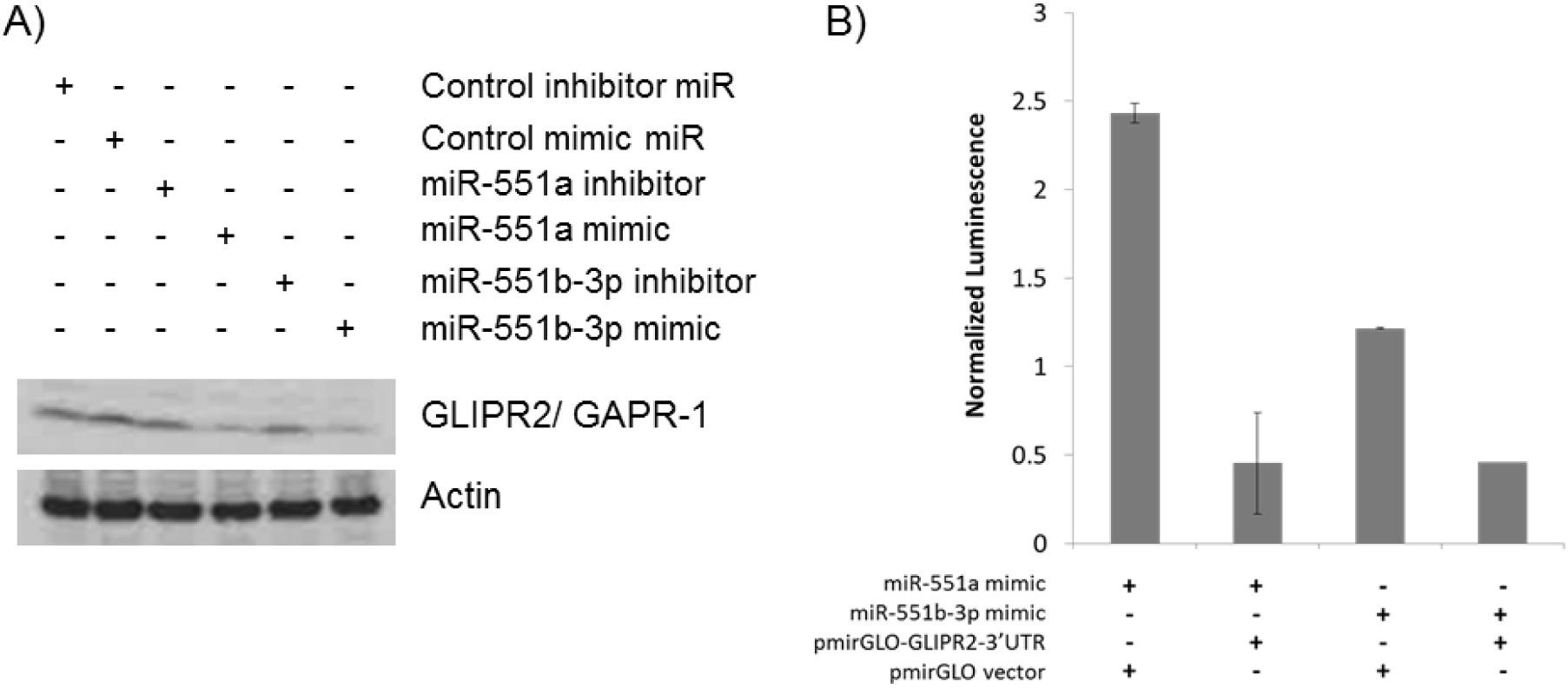
miR-551a and miR-551b-3p target GLIPR2. A) Transfection of miR-551a and miR-551b-3p mimics resulted in decreased expression of GLIPR2 where as their inhibitors have showed almost no effect on GLIPR2 expression. No effect upon inhibitors treatment could be due to saturated levels of GLIPR2 expression or not enough concentration of inhibitors in HN5 cells. B) Luciferase reporter assay using 3’UTR region of GLIPR2 showing that miR-551a and miR-551b-3p physically interacts with 3’UTR region of GLIPR2 and their physical interaction upon transfection of miRNAs mimics resulted in decreased luciferase activity indicating that miR-551a and miR-551b-3p directly interacts with GLIPR2 3’UTR region to target its expression. cDNA transfection at different concentrations as shown in above leads to reduced proliferation which is agreement with miR-551a and miR-551b-3p inhibitor treatment condition in HN5 cells.

### GLIPR2 has negative effect on HN cancer cell survival, migration and invasion

GLIPR2 cDNA transfection into HN5 and UMSCC-17B resulted in significant decrease in cell survival (Fig 8.A). We observed GLIPR2 cDNA concentration dependent reduction in cell proliferation using CTG assay. Besides cell proliferation GLIPR2 cDNA decreased migration (Fig 8.B) and invasion (Fig 8.C) also in HN cancer cells.

**Fig.8:**
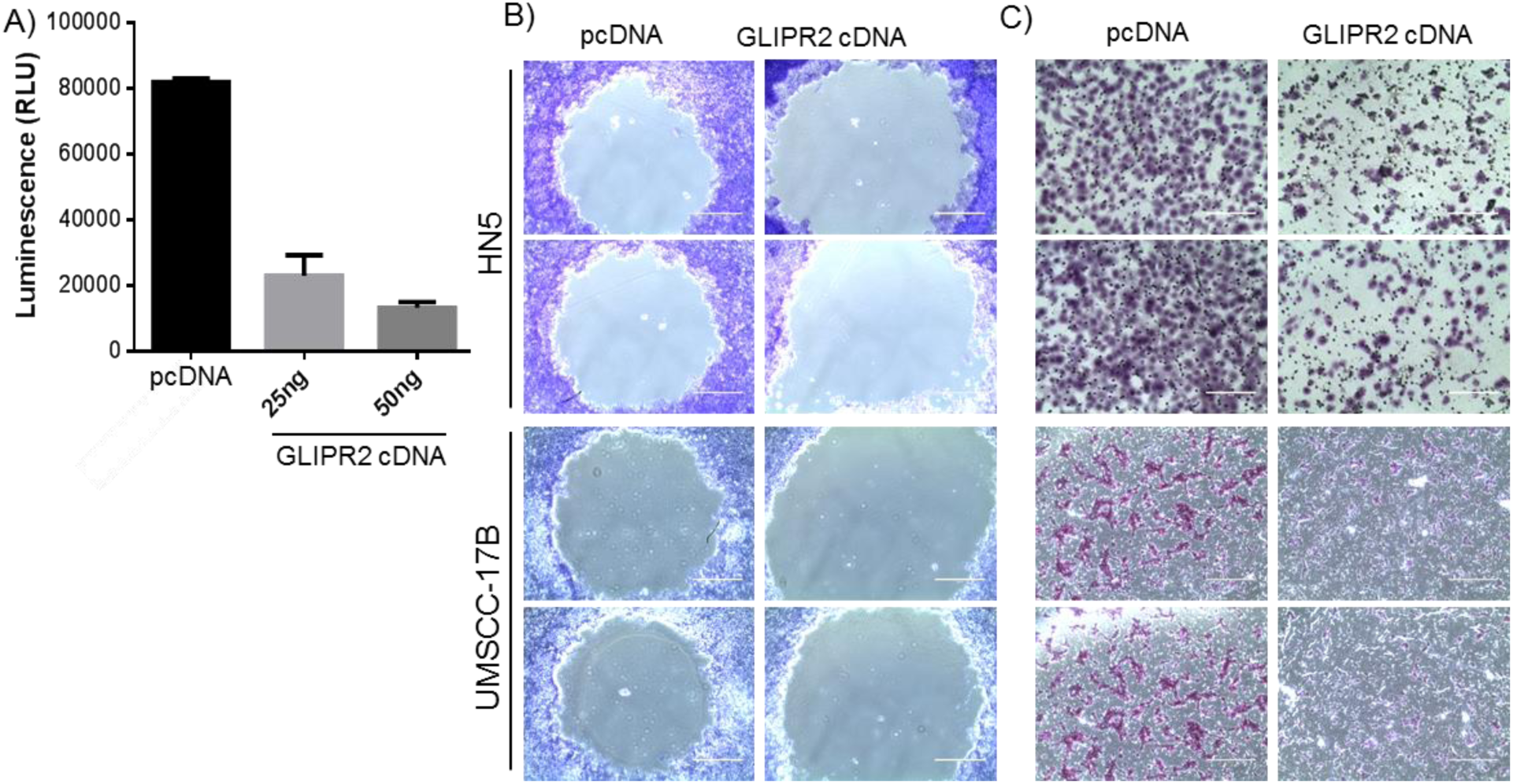
GLIPR2 inhibits cell proliferation and invasion and increases radio sensitization. Transfection of GLIPR2 cDNA resulted in decreased A) cell proliferation and B) invasion, which is agreement with miR-551a and miR-551b-3p inhibitor treatment condition in HN5 cells. C) Ectopic expression of GLIPR2 cDNA decrease cell survival fraction in HN5 cells upon radiation compared to control pcDNA. Survival fraction curves were drawn using linear quadratic fit method using Graphpad Prism.

### miR-551a and miR-551b-3p increases systemic autophagy and activates PI3-AKT and MAPK pathway

miR-551a and miR-551b-3p mimics transfection into HN cancer cells increased the systemic autophagy which was evident by increased LC3-I to LC3-II conversion and this was reversed by miR-551a and miR-551b-3p inhibitors. Interestingly p62 protein levels were accumulated in miR-551a and miR-551b-3p mimics transfected condition (Fig 9.B). This result was reiterated by using GLIPR2 siRNA transfection which indirectly reflects miR-551a and miR-551b-3p mimics transfection condition. Interestingly miR-551a and miR-551b-3p mimics transfection increased AKT phosphorylation and p38 MAPK phosphorylation as well while decreasing p27-kip1 levels (Fig 9. C) which drive the cells into active proliferation, migration and invasion direction.

**Fig.9:**
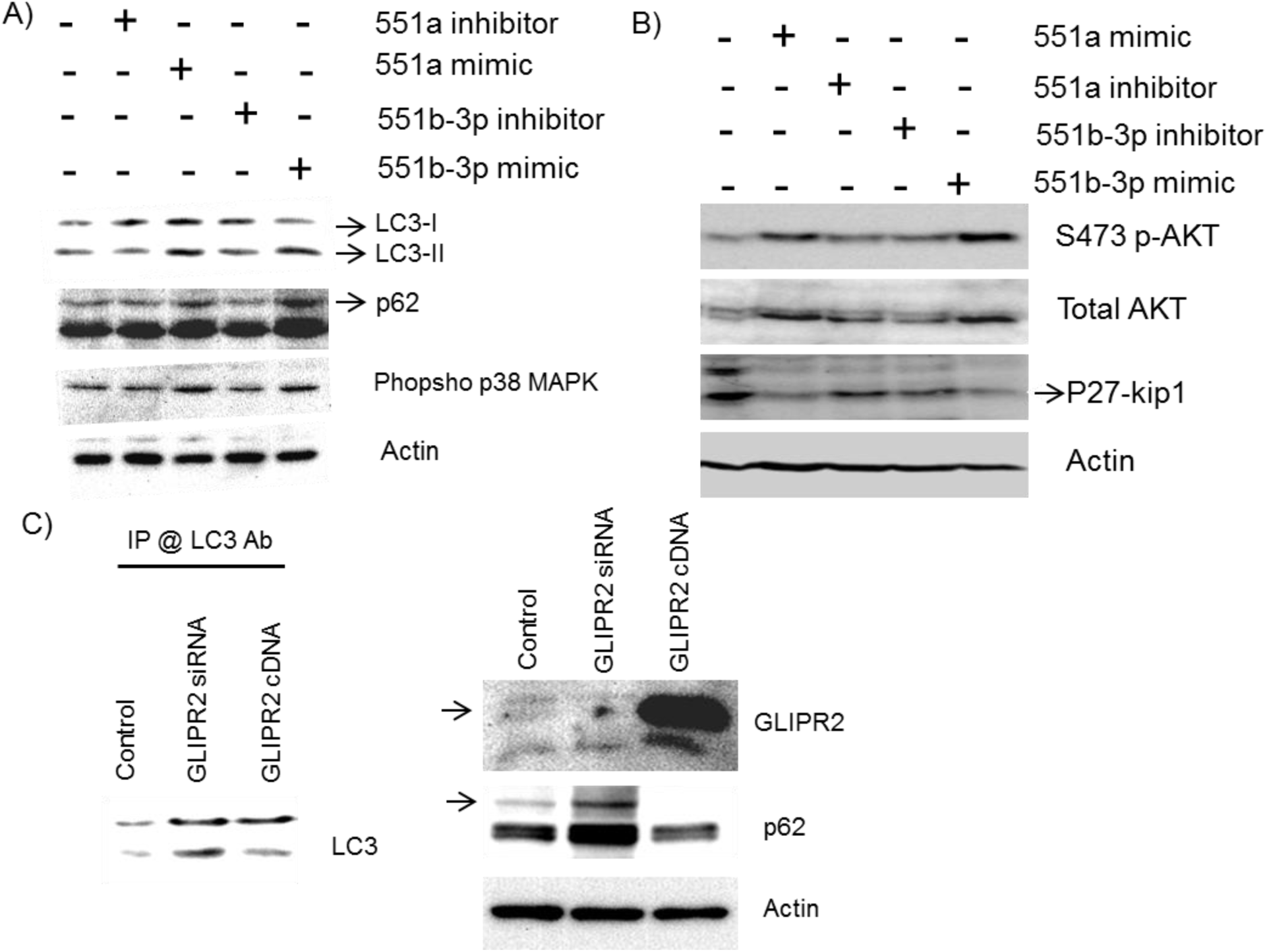
miR-551a and miR-551b-3p targeted GLIPR2 axis involved in increased and sustained autophagy there by activation of p38 MAPK and AKT pathway. A) Transfection of miR-551a and miR-551b-3p mimics induces autophagy where as their inhibitor decreases which is illustrated by increased/decreased LC3-II intensity and associated with p62 protein levels and phospho p38 MAPK levels as well. B) Transfection of miR-551a and miR-551b-3p mimics increases phosphorylation of AKT and decreases p27 kip1 expression. C) Ectopic expression of GLIPR2 cDNA decreases and transfection of GLIPR2 siRNA increases autophagy and it correlated with p62 protein levels also.

### GLIPR2 expression value correlates with stage specific patients overall survival

Upon understanding the function of and GLIPR2 we tried to find out the relation between GLIPR2 expression and survival in HN cancer patients in our dataset and TCGA data set as we did in case of miR-551a, miR-551b-3p expression. We found that though the trend is in an expected way, p value is not significant when total patients’ population was divided based on median value of GLIPR2 expression values across the all patients. Interestingly GLIPR2 expression alone was not significantly associated with survival of patients though it is in expected trend but GLIPR2 to Beclin1 ratio was significantly associated with overall survival of HN cancer patients in TCGA data set (Fig.10). Recently it was found that Beclin1 interacts with GLIPR2 which is a negative regulator of autophagy (Shoji-Kawata et al., 2013a). So we postulate that GLIPR2 expression with respect to Blecin1 is more clinically relevant than these two individual proteins expressions alone.

**Fig.10:**
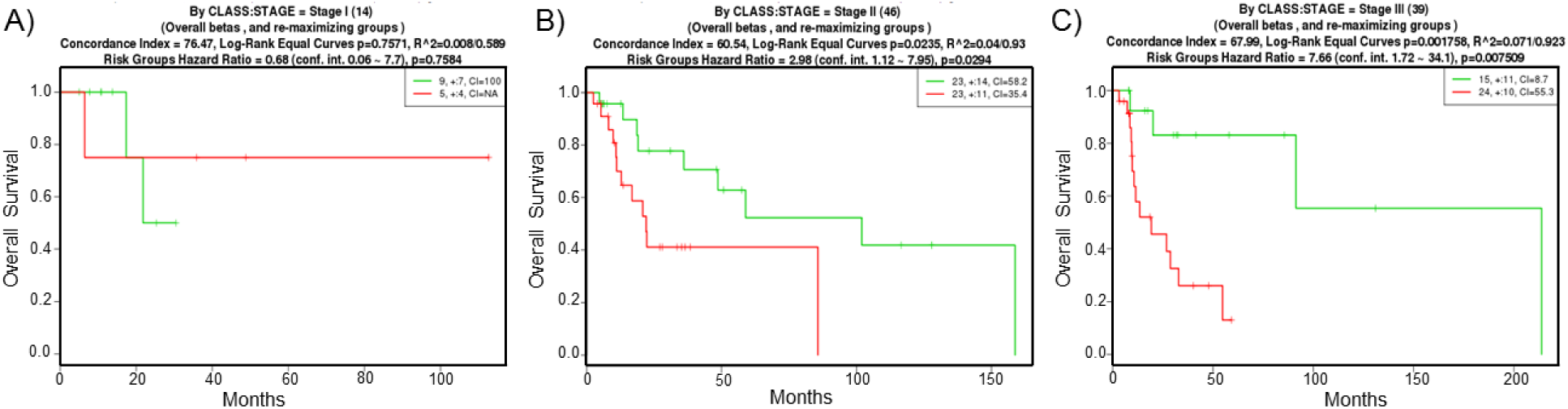
GLIPR2 expression value correlates with stage specific patients overall survival. In TCGA data set patients were divided based on their HN cancer stage and GLIPR2 expression values correlate with overall survival of C) Stage III patients with high p-value of 0.007 while it is less significant in B) Stage II with p-value of 0.029 and A) Stage I with p value of 0.75.

### miR-551a and miR-551b-3p target GLIPR2 regulates proliferation through autophagy

To verify whether GLIPR2 induced autophagy inhibition is a potential mechanism underlying the observed increased proliferation by miR-551a and miR-551b-3p at least in part we have conducted proliferation experiments as described earlier in presence and absence of Bafilomycin A1 which is a potent autophagy inhibitor. We transfected two sets of HN5 cells with miR-551a, miR-551b-3p mimics and inhibitors and GLIPR2 siRNA, cDNA and then one set was incubated with 100nM BafilomycinA1 and another set was used as control. We observed increased proliferation when transfected with miR-551a and miR-551b-3p mimics and decreased proliferation wen transfected with miR-551a and miR-551b-3p inhibitors and GLIPR2 siRNA as shown earlier. Interestingly we observed such effects were almost mitigated to control level in presence of Bafilomycin A1 suggesting miR-551a, miR-551b-3p and GLIPR2 mediated autophagy role in cell proliferation (Fig.11).

**Fig.11:**
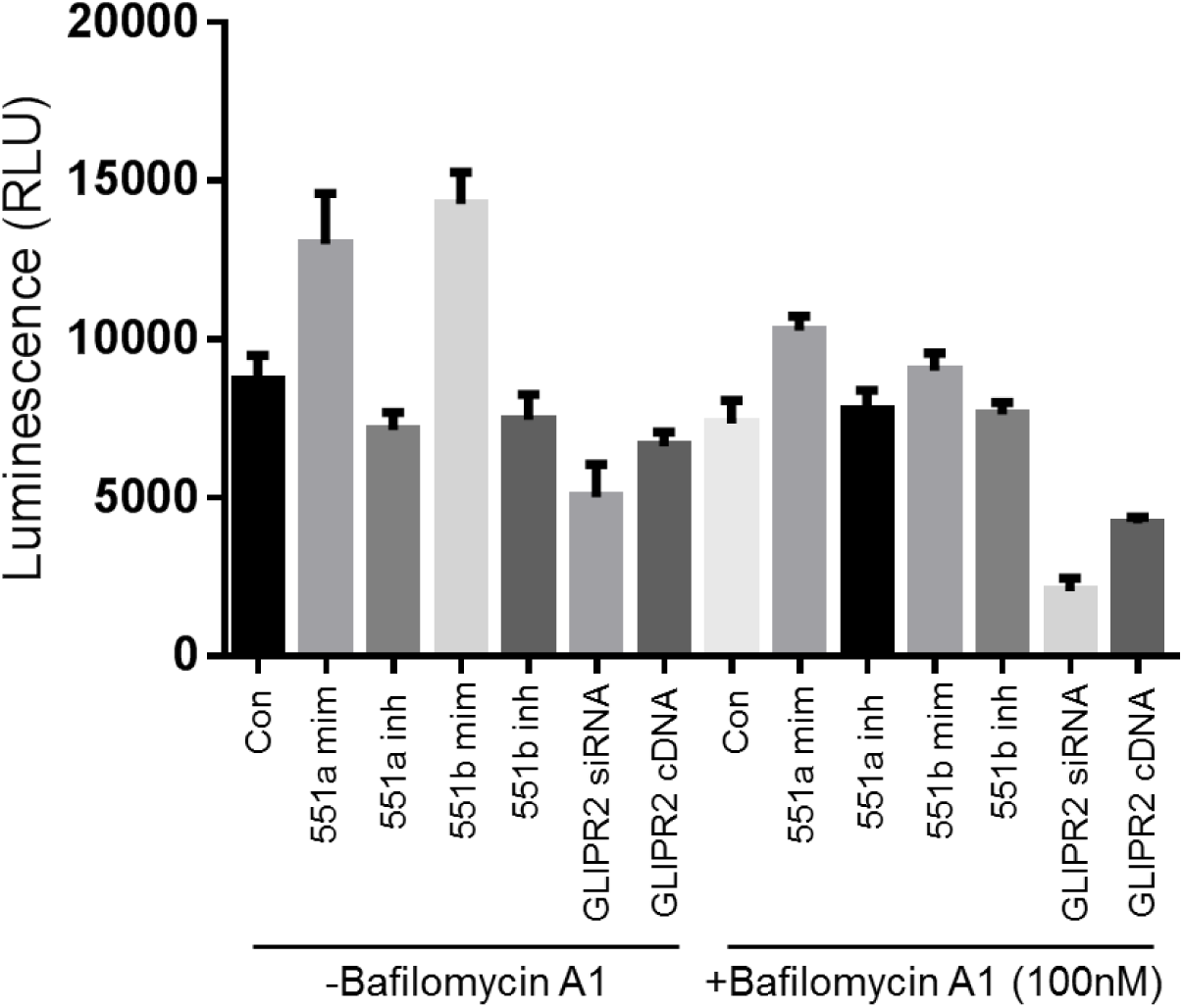
Autophagy inhibitor Bafilomycin A1 nullifies the pro-proliferative effect of miR-551a, miR-551b-3p. In presence of 100nM Bafilomycin A1 increased proliferation by miR-551a, miR-551b-3p mimics and decreased proliferation by miR-551a, miR-551b-3p inhibitors was nullified to almost control level.

## Discussion

Personalized medicine is now valued as one of the successful approach to improve the survival and quality of life of patients especially who are suffering advanced and high grade tumors. (Bellmunt and Petrylak, 2012; Rahman et al., 2012; Van Allen and Pomerantz, 2012). In order to move in the direction of personalized medicine one need to understand the molecular mechanisms underlying tumorigenesis and progression. Towards understanding molecular nature of HNSCC carcinogenesis and its progression we have conducted miRNA profiling experiment of xx number of HN tumor samples. Most of these patients have undergone post-operative radio therapy (PORT) including chemo therapy (details) and found to be at high-risk for recurrence and secondary tumors. There are 28 number of differentially expressed miRNAs from the study which includes already known to be involved in carcinogenesis and tumor progression. From this list of differentially expressed miRNAs we have picked miR-551b-3p which holds very good p value of 0.04 and ∼1.5 fold change between DM and NED patients. qRT-PCR results suggested that there is about ∼2.1 fold increased expression of miR-551b-3p in DM patients compared to NEDs. This increase in fold change by qRT-PCR results could be due to the fact that fold change measures on miRNA array platform is more compressed (Pradervand et al., 2009). To our knowledge there are no reports so far on miR-551b-3p role in tumorigenesis and tumor progression. miR-551b-3p is localized on chromosome 3: 168551854-168551949 of positive strand (miRBase.org). Interestingly this region was found to be amplified in HN tumors of our data set by aCGH analysis (data not shown). Later on we found that miR-551a also contains exactly same seed sequence as in miR-551b-3p and miR-551a also up regulated moderately (∼1.2 fold) in DM patients compared to NEDs with p value of 0.04. miR-551a is present on chromosome 1:3560695-3560790 on negative strand. We hypothesize that both these miRNAs could exhibit similar biological effects by the virtue of having same seed sequence, however their spatial and temporal and conditional dependent expression regulation might be important in deciding their function and cell phenotype. So far there are two reports about miR-551a in connection with cancer progression. Earlier Li Z et al found that miR-551a and miR-495 target Phosphatase of regenerating liver-3 (*PRL-3*) there by act as tumor suppressor in human gastric cancer (Li et al., 2012). Very recently Loo JM et al reported that miR-551a together with miR-483 target creatine kinase brain type (CKB) and suppress colorectal cancer progression (Loo et al., 2015). However we found that miR-551a and miR-551b-3p upregulated in HN tumors and validated by qRT-PCR. This difference in direction of miR-551a expression could be due to the fact that we are studying miR-551a role in secondary HN tumors and earlier reports are in primary tumors of colorectal and gastric cancers. There are some earlier reports showing this kind of opposite expression changes in terms of genes and miRNAs in different kinds of tumors (Volinia et al., 2006). This kind of discrepancy could be mainly because of different cancer cell origin or stage of tumors.

Our observation about miR-551a and miR-551b-3p up-regulation in HN patients with DM compared to NED after PORT was well justified by the Kaplan-Meir survival analysis using the patients overall survival data after surgery. In our data set DM patients containing higher expression of miR-551a and miR-551b-3p have lower survival probability compared to NED patients containing low expression of miR-551a and miR-551b-3p (Fig.2). In addition to our data set we also found the same correlation between miR-551a and miR-551b-3p expression levels and survival probability in another nasopharyngeal carcinoma data set (GSE36682) (Antonov, 2011; Antonov et al., 2013) which is one of the HN tumor type.

Tumor progression is mainly derived by three salient features of cancer cells such as proliferation, migration within the organ and invasion to other parts of human body (Limame et al., 2012). Hence we have tested the effect of miR-551a and miR-551b-3p on these features using synthetic mimics and inhibitors. From these experiments we found that miR-551a and miR-551b-3p mimics indeed have positive effect on proliferation, migration and invasion capabilities of HN cancer cells in-vitro and their inhibitors exerted negative effect. Hence our observation from miRNA array and qRT-PCR results that increased miR-551a and miR-551b-3p expression in patients who had poor prognosis and distant metastasis is in full accordance with their newly unraveled functional role in HN cancer cells. We even found that miR-551a and miR-551b-3p mimics induce AKT and p38 MAPK phosphorylation while suppressing p27 kip1 which could explain the signaling pathways involved in promoting proliferation, migration and invasion (Matsuoka and Yashiro, 2014).

Generally miRNAs exerts their functions through modulating its target mRNA expression by binding to its 3’UTR of target mRNA (Carninci et al., 2005). There are several target prediction softwares’ available now to know certain miRNA’s putative targets. But we took an approach to evaluate miR-551a and miR-551b-3p target by conducting micro array experiment using total RNA extracted from HN5 cells transfected with miR-551a and miR-551b-3p mimics and inhibitors. We focused only one those mRNAs who had decreased expression in miR-551a and miR-551b-3p mimics treated condition whereas an increased expression in miR-551a and miR-551b-3p inhibitors treated condition compared to their respected controls. This approach led us to the discovery of GLIPR2 as potential target of miR-551a and miR-551b-3p both and then we checked using MIRSYSTEM (http://mirsystem.cgm.ntu.edu.tw/) (Lu et al., 2012) that what is the exact sequence in 3’UTR of GLIPR2 which is complementary to miR-551a and miR-551b-3p seed sequence. Using luciferase reporter assay and westernblot experiment containing this sequence we confirmed that GLIPR2 is a target of miR-551a and miR-551b-3p. Interestingly GLIPR2 had inverse correlation in terms of expression compared to miR-551a and miR-551b-3p expression in our data set i.e. DM patients had significantly lower expression of GLIPR2 compared to NED patients.

GLIoma Pathogenesis Related protein2 (GLIPR2) is a small protein of 17.2 kDa which contains 154 amino acids belongs to the super family of plant pathogenesis related proteins (Eberle et al., 2002). GLIPR2 was first identified to be localized to lipid-enriched microdomains in the Golg complex of mammalian cells hence also known as Golgi associated plant pathogenesis related protein-1 (GAPR1). GLIPR-2 protein known to be N-myristoylated and predominantly expressed in monocytes, the lung, spleen, kidney lymphocyte, uterus and embryonic tissue. GLIPR2 was found to dimerize and form and active-site across the dimer interface which contains putative catalytic residues for serine protease activity (Serrano et al., 2004). GLIPR2 was reported to be highly expressed iin fibrotic kidney and promotes epithelial to mesenchymal transition (EMT) in vitro (Baxter et al., 2007). Recently it was reported that EMT of hepatocellular carcinoma cells was promoted by hypoxia induced GLIPR2 expression (Huang et al., 2013). However from micro array gene expression data we observed decreased expression of GLIPR2 mRNA expression in DM patients which are likely to have more hypoxic micro environment. Another report stated that GLIPR2 overexpression promotes EMT and migration through activation of ERK1/2 signaling pathway in human proximal renal tubular epithelial (HK-2) cells (Huang et al., 2013). In our study we found that GLIPR2 cDNA transfection reduces proliferation, migration and invasion in HN5 and UMSCC-17B HN cancer cell lines. These results resemble the situation of miR-551a and miR-551b-3p mimics transfecting condition.

Another recent report by Shoji-Kawata et al, suggested an important autophagy negative regulatory role of GLIPR2. They found that GLIPR2 is Beclin 1 interacting protein (Shoji-Kawata et al., 2013b). Autophagy is a mechanism by which cellular homeostasis is maintained through selective degradation of cellular components including long-lived proteins, protein aggregates, damaged cytoplasmic organelles which in turn recycles the nutrients and energy generation (Levine and Klionsky, 2004). However autophagy was shown to have both tumor suppressor and oncogenic roles depending on the context. While excessive autophagy leads to cell death, under stress conditions autophagy promotes cell death (Puissant et al., 2012). Here in our study we found that upon miR-551a and miR-551b-3p transfection an increase in autophagy which is denoted by increased LC3-I to LC3-II conversion. Generally with increasing autophagy p62 is degraded which serves as molecular adaptor between autophagic machinery and its substrates. Interestingly, we observed accumulation of p62 protein also in these samples. Recently it was shown that p62 accumulation promotes proliferation, migration and EMT process through stabilization of TWIST1 protein (Qiang et al., 2014). We assume that this same scenario was replicated using GLIPR2 siRNA also. These results suggests that there is autopagy defect in these HN cell lines which is enhanced by miR-551a and miR-551b-3p and this accumulation of p62 which can dissociate NRF2 from KEAP1 which induces anti-oxidant response pathway by binding to antioxidant regulatory element motif (Itoh et al., 1999; Jain et al., 2010; Komatsu et al., 2010). We even observed increase in NRF2 target genes such as NQO1, GSTA1, ATF3 etc., up regulation in DM patients compared to NED patients (Fig.12). These NRF2 target genes were earlier reported by CHIP-Seq experiments (Chorley et al., 2012).

**Fig.12:**
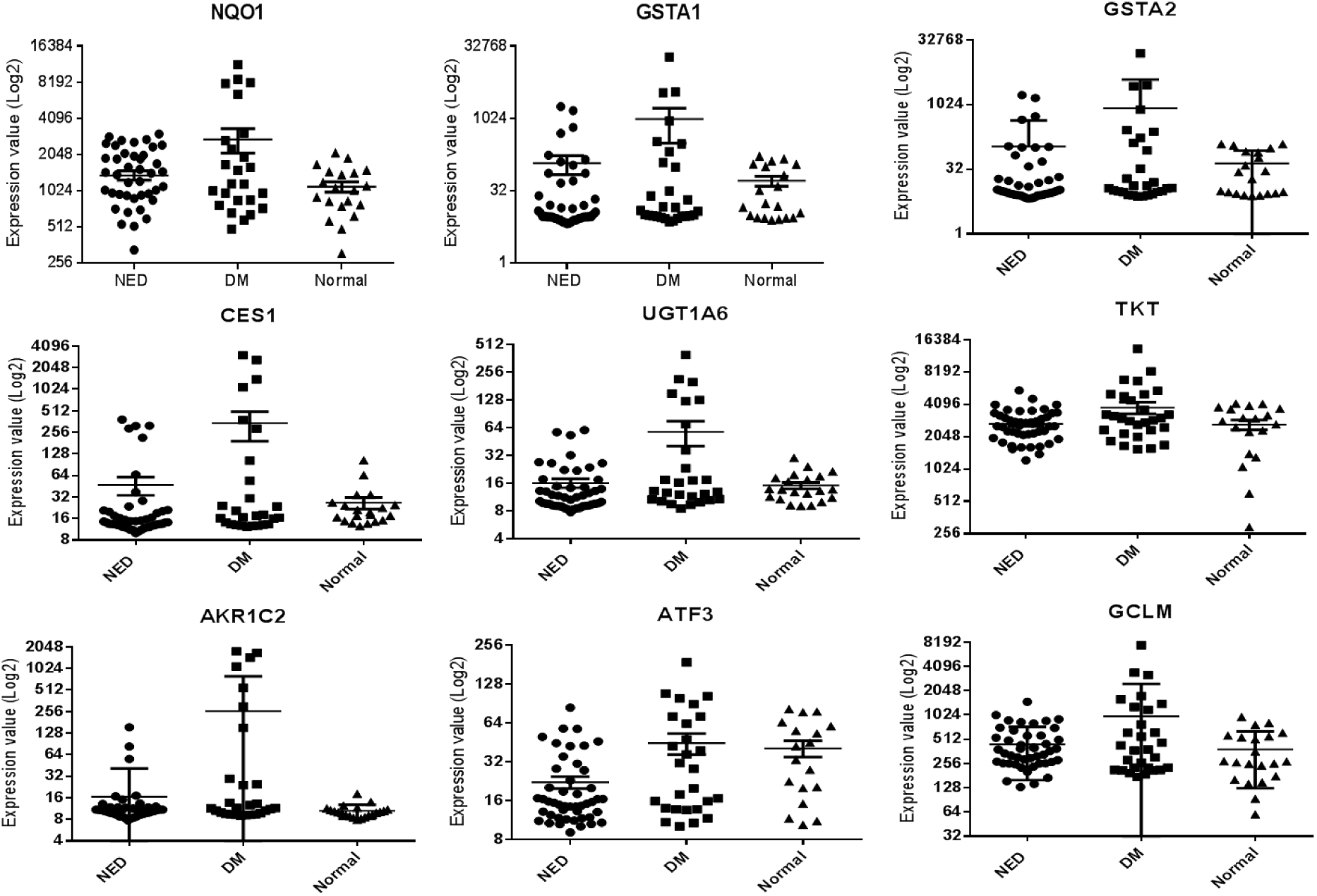
Gene expression values of NRF2 target genes in UTSW HN tumor data set. Log2 gene expression values graphs were drawn using HN cancer patients’ gene expression array results for each condition in our data set for NQO1, GSTA1, GSTA2, CES1, UGTA6, TKT, AKR1C2, ATF3 and GCLM genes which are known NRF2 targets.

GLIPR2 expression is positively correlated with survival of patients in our data set with p value of 0.08. This could be because the main function of GLIPR2 might be associated with Beclin-1 interaction which is main effector of autophagy. To support this notion we found that GLIPR2/Beclin1 ratio better correlate (p value= 0.04) with survival probability in TCGA HN cancer data set than GLIPR2 (p value= 0.12) or Beclin1 (p value= 0.06) individually. It was very interesting to observe GLIPR2 expression value association with higher stage HN tumor patients’ survival in TCGA data set (Fig.10). Though GLIPR2 expression is positively correlated with survival of all stages of patients in TCGA data set it was not significant with p value of 0.09. But when these patients were classified according to their stages of cancer then the correlation between GLIPR2 expression and patients’ survival was highly significant in Stage III patients (p value=0.007) while in Stage II it was significant with p value of 0.029 and in Stage I it was not significant with p value of 0.75. This result suggests that miR-551a, miR-551b-3p and GLIPR2 axis has more significant impact in higher grade tumors compared to lower grade tumors.

Taken our all results together we propose that miR-551a and miR-551b-3p function as oncogenic micro RNAs in at least HN cancer setting and they target GLIPR2. miR-551a and miR-551b-3p exerts their function at least in part through GLIPR2 by modulating autophagy in HN cancer cells and activating PI3-AKT, p38 MAPK signaling pathway.

## Acknowledgements

We thank Suneetha Reddy Aluru for her technical support. This work was supported by Cancer Prevention and Research Institute of Texas (CPRIT-5000764303) funding to MDS.

